# Pleistocene climate changes explain large-scale genetic variation in a dominant grassland species, *Lolium perenne* L

**DOI:** 10.1101/414227

**Authors:** J.L. Blanco-Pastor, S. Manel, P. Barre, A.M. Roschanski, E. Willner, K.J. Dehmer, M. Hegarty, H. Muylle, T. Ruttink, I. Roldán-Ruiz, T. Ledauphin, A. Escobar-Gutierrez, J.P. Sampoux

**Author notes:** **Corresponding author:** J.L. Blanco-Pastor, INRA, Centre Nouvelle-Aquitaine-Poitiers, UR4 (URP3F), Le Chêne - RD 150 CS 80006, 86600 Lusignan, France.

## Abstract

**Aim:** Grasslands have been pivotal in the development of herbivore breeding since the Neolithic and are still nowadays the most widespread agricultural land-use across Europe. However, it remains unclear whether the current large-scale genetic variation of plant species found in natural grasslands of Europe is the result of human activities or natural processes.

**Location:** Europe.

**Taxon:** *Lolium perenne* L (perennial ryegrass).

**Methods:** We reconstructed the phylogeographic history of *L. perenne*, a dominant grassland species, using 481 natural populations including 11 populations from closely related taxa. We combined the Genotyping-by-Sequencing (GBS) and Pool-sequencing (Pool-seq) methods to obtain high-quality allele frequency calls of ~ 500 k SNP loci. We performed genetic structure analyses and demographic reconstructions based on the site frequency spectrum (SFS). We additionally used the same genotyping protocol to assess the genomic diversity of a set of 32 cultivars representative of the *L. perenne* cultivars widely used for forage purposes.

**Results:** Expansion across Europe took place during the Würm glaciation (12-110 kya), a cooling period that decreased the dominance of trees in favour of grasses. Splits and admixtures in *L. perenne* fit historical sea level changes in the Mediterranean basin. The development of agriculture in Europe (7-3.5 kya), that caused an increase in the abundance of grasslands, did not have an effect on the demographic patterns of *L. perenne*. We found little differentiation between modern cultivars and certain natural variants. However, modern cultivars do not represent the wide genetic variation found in natural populations.

**Main conclusions:** Demographic events in *L. perenne* can be explained by the changing climatic conditions during the Pleistocene. Natural populations maintain a wide genomic variability at continental scale that has been underused by recent breeding activities. This variability constitutes valuable standing genetic variation for future adaptation of grasslands to climate change, safeguarding the agricultural services they provide.

## Introduction

Worldwide, grasslands constitute the most extensive natural and semi-natural habitat types and are an integral part of agricultural landscapes. They have provided ecological services that have been essential for the maintenance of biodiversity, carbon sequestration and the biogeochemistry of soils across the last millennia (Tilman et al., 1996; Jones & Donnelly, 2004; Hejcman et al., 2013). Grasslands, have supported biodiversity and a variety of ecosystem services that have gained renewed interest and value to society nowadays (Werling et al., 2014). They are furthermore the most widespread agricultural land-use in Europe, covering 45% of the total crop area in the European Union (Eurostat, 2017). Most of these grasslands are permanent natural or semi-natural grasslands (84%; Leclère et al., 2016) and are composed of plant species and populations that have evolved under both environmental conditions and farming usages at least since the earliest stages of herbivore domestication and agriculture, i.e. about 10 kya in the Fertile Crescent (Hejcman et al., 2013). Conversely, temporary grasslands (or meadows) (14%; Leclère et al., 2016) are sown for a period of one to five years and are intensive herbivore production systems mainly composed by modern cultivars. Since years 1960, experimental agronomic research centres have implemented breeding programs to develop improved cultivars ensuring high production of forage of good quality (Sampoux et al., 2011). New permanent or temporary grassland acreages are nowadays almost exclusively sown with such cultivars.

The maintenance of plant genetic diversity in grassland species can be of major importance not only in terms of adaptive potential but also in terms of productivity and ecosystem services (Crutsinger et al., 2006; Hughes et al., 2008). It has been shown that intra-population genetic diversity may favour the temporal stability of production in grasslands (Prieto et al., 2015). Also large-scale genetic variation may be tightly linked with regional productivity of grasslands dominated by perennial ryegrass (Balfourier & Charmet, 1991; Monestiez et al., 1994). Since the development of modern plant breeding, the delivery of improved forage grass cultivars has led to a rapid expansion of the acreage of sown meadows whereas the acreage of natural grasslands has continuously decreased (Chapman, 1992). In the EU-6 (first six members of the European Union), losses of permanent grasslands are estimated at about 30% (7 million ha) between 1967 and 2007 whereas same losses in EU-15 are estimated at 15% (10.5 million ha) during the last 50 years (Peeters, 2012). Given this trend, it remains particularly relevant to assess the risk of genetic impoverishment of grassland species across agricultural landscapes that may lead to losses of adaptive potential and/or production capability.

Reconstructions from the pollen record revealed the extensive presence of open habitat (*i.e.* grasslands) in Europe since the pre-Holocene (> 12 ka ago –kya) (Pèrez-Obiol & Julià, 1994; Kuneš et al., 2015). Grasslands in central-eastern Europe, for example, constituted up to 30% of the vegetation already during the early-Holocene and their relative abundance has prevailed until current time (Kuneš et al., 2015). Despite the fact that grasslands were widespread long before agricultural practices became established in Europe *ca.* 8 kya (Zohary et al., 1988; Hejcman et al., 2013; Giesecke et al., 2017), activities of early agricultural communities have been considered to be important for shaping current European vegetation and maintaining open land (Feurdean et al., 2015). Pollen records clearly show that the area dominated by grasses in Europe increased since 4 kya, suggesting that the conversion of forest landscapes into grasslands was associated with the development of agriculture (Giesecke et al., 2017). However, it is not clearly known whether this human transformation of the European landscape also affected the natural diversity and population structure of grassland species. More specifically, it is unknown the relative importance of natural versus human-mediated expansion of grasses to explain the current genetic variation in natural and semi-natural grasslands. Additionally, the risk of genetic diversity loss in grassland species due to the increase use of cultivars at the expense of natural grasslands remains unevaluated. To investigate these questions we reconstructed the phylogeography of *Lolium perenne* L. (perennial ryegrass) and assessed its current genetic variation at a continental scale. We used *L. perenne* as a model to represent the genetic diversity in European grasslands because this species is the most widely sown forage grass species in temperate regions (Humphreys et al., 2010), has received extensive breeding effort during the last four decades (Humphreys et al., 2010; Sampoux et al., 2011) and is also one of the most abundant grass species in natural grasslands across Europe and the Fertile Crescent.

Based on chloroplast DNA (cpDNA) polymorphisms, it was suggested that natural populations *L. perenne* could have been subjected to unconscious selection since the Neolithic and introduced to Europe following human migration routes from the Fertile Crescent as a weed of cereals (Balfourier et al., 2000). More recently, a study based on the analysis of 2185 transcript-anchored SNPs has suggested that *L. perenne* was subjected to repeated population expansion and contraction during the Quaternary glaciations (Blackmore et al., 2015). However, the latter study did not estimate the time of demographic events, which is required to firmly discard the hypothesis of human-mediated expansion. One complicating factor for the study of *L. perenne* is the fact that the genus *Lolium* comprises nine species (Terrell, 1968; Charmet et al., 1996) that diverged only recently (ca. 4.1 Ma ago; Inda et al., 2014) and that can intercross among themselves. This explains the controversial phylogenetic relationships between *L. perenne* and close relatives observed in the literature (Charmet et al., 1997; Catalán et al., 2004; Inda et al., 2014).

Thanks to the advances in sequencing technologies (Emerson et al., 2010; Garrick et al., 2015), assessing fine-scale phylogenetic, phylogeographic and hybridization patterns among recently diverged lineages is now an achievable task. Furthermore, recently developed methods that sequence pools of individuals to represent the genomic constitution of a population (Pool-seq; Schlotterer et al., 2014), especially in combination with reduced representation libraries such as Genotyping-by-Sequencing (Byrne et al., 2013), have opened opportunities for the genome-wide genotyping of numerous populations at affordable cost. These advances combined with novel statistical models based on site frequency spectrum analysis (Gutenkunst et al., 2009; Excoffier et al., 2013) make now possible the assessment of past demographic dynamics in closely related lineages with unprecedented detail. This enables the estimation of structure and past demography even in species such as *L. perenne*, with a continuous distribution across wide geographical regions and exhibiting high heterozygosity levels due to a very effective self-incompatibility system that promotes outbreeding (Cornish et al., 1979).

Here we present a comprehensive study of the genomic diversity of natural *L. perenne* populations and reconstruct historical demographic events that contributed to shape this diversity, using an extensive set of populations sampled in natural and semi-natural grasslands across almost all the entire natural distribution range of the species. In order to capture the broadest possible spectrum of natural lineages within *L. perenne*, we sampled 470 natural populations across Europe and the Fertile Crescent. This panel was complemented by several populations from other closely related taxa, namely five populations from *L. multiflorum*, two from *L. rigidum*, two from *L. temulentum* and two from *Festuca pratensis*. Additionally, we selected a set of 32 *L. perenne* cultivars representing the cultivated gene pool bred for forage usage in Europe and New Zealand. All populations were genotyped with GBS Pool-seq, yielding a filtered set of ca. 500 k SNP loci, randomly distributed across the genome. Specific objectives of this study were: (i) to reveal genetic structure, gene flow and trace historical demography from the natural diversity of *L. perenne*; (ii) to investigate the relationships between *L. perenne* and close relatives; and (iii) to get insight into the origin of *L. perenne* cultivars and trace the use of natural genetic resources in this species by modern plant breeding. A better knowledge of the distribution of the natural genomic variation of *L. perenne* across its area of primary expansion in Europe would contribute to the understanding of factors that shaped the genetic diversity of grass species in current natural and semi-natural grasslands. It would also provide essential information to guide the conservation of the diversity of this major grassland species in a context of reduction of the acreage of permanent natural grasslands. It would also guide the informed use of natural populations as genetic resources for plant breeding.

## Materials and Methods

### Plant Material and Genotyping

Detailed information on the origin of the plant material used in the present study, the genotyping and the genomic data filtering can be found in the Supporting Information Methods S1.

### Phylogeography and past demography

#### Population structure

We used the Discriminant Analysis of Principal Components (DAPC, Jombart *et al.*, 2010) analysis on the *L. perenne* set, a clustering method that does not rely on a particular population genetics model. DAPC was conducted with the R package ‘adegenet’ (Jombart, 2008; Jombart & Ahmed, 2011). We first applied the k-means procedure implemented in the function *find.cluster* to infer the optimal number of clusters *K* (Jombart et al., 2010). *K* was determined using the Bayesian Information Criterion (BIC). The function BIC = *f*(*K*) was U-shaped, so *K* was selected as the value after which the decrease of BIC was much less steep (Fig. S1), as recommended in Jombart et *al.* (Jombart et al., 2010). We used 10,000 iterations of the K-means algorithm and 100 randomly chosen starting centroids in each run to ensure convergence of the algorithm. The number of Principal Components (PC) retained for the discriminant analysis is a trade-off between power and consistency of assignment. We therefore determined the optimal number of components to be retained by calculating the mean a-score from ten runs of the DAPC (optim.a.score function in adegenet). We also computed Fst among the clusters obtained by the k-means algorithm with the R package ‘StAMPP’ (Pembleton et al., 2013). Finally, we tested isolation by distance (IBD) using a Mantel test (Mantel, 1967) with the R packages ‘geosphere’ (Hijmans et al., 2012) and ‘vegan’ (Oksanen et al., 2011). Additionally, we calculated the expected heterozygosity (*He*) of populations and computed the *He* average value and standard deviation of each cluster. DAPC analyses were similarly carried out on the 79 *cpDNA* SNP loci (*L. perenne* set and *Lolium* set).

#### Splits and admixture

We analysed the nuclear genomic data with the program TREEMIX v.1.13 (Pickrell & Pritchard, 2012). Before running TREEMIX in our dataset, allele frequency data were averaged over populations from a same cluster, as defined by DAPC. We ran two TREEMIX analyses with different datasets. The first analysis was applied to a dataset including only data from *L. perenne* clusters (*L. perenne set*). This dataset contained 507,583 SNP loci free of missing data. The second analysis was applied to a dataset including data from *L. perenne* clusters (470 populations) together with outgroup populations from *F. pratensis* (2), *L. multiflorum* (5), *L. temulentum* (2) and *L. rigidum* (2) (*Lolium set*). To avoid the presence of early generation hybrids in the outgroup populations, we only included those populations unambiguously determined with on-field taxonomy observations. This second dataset contained 131,113 SNP loci free of missing data. For both datasets, we first inferred a maximum likelihood (ML) tree without admixture in TREEMIX and then ran 1000 standard bootstrap replicates to obtain statistical support for the non-admixed tree topology. Bootstrap trees were summarized in a 50% majority-rule consensus tree with SumTrees v4.2.0 (Sukumaran & Holder, 2015). Finally, for both datasets we also fitted one to six migration events in the tree and displayed a tree graph for each number of migration events.

#### Demography

To investigate the hybridization patterns among the *Lolium* taxa included in our study and the demographic history of *L. perenne*, we used the software *δaδi* (Gutenkunst et al., 2009). *δaδi* is a method that compares the SFS expected under custom demographic models to that observed with our data. SFSs are simulated with a diffusion approach which is limited to a three-taxon phylogeny as implemented in *δaδi*. Comparisons of *δaδi* models were made independently for each of the two datasets (*L. perenne* and *Lolium*). For the construction of *δaδi* models we used alternative migration/hybridization scenarios as obtained with TREEMIX. First, we investigated demography and patterns of gene flow within *L. perenne,* as displayed in the *L. perenne set* TREEMIX models (TMMs) 1-6 (see below). For this analysis, we averaged frequency data of populations from clusters 1 to 5 and considered clusters 6 and 7 as independent lineages (as displayed in the main topology of *L. perenne set* TMMs 1-6, see below). Second, we investigated alternative scenarios of gene flow among *L. rigidum, L. multiflorum* and *L. perenne* as shown in the *Lolium set* TMMs 1-6 (see below). For this second analysis, we averaged frequency data of all populations from *L. perenne* genetic clusters. For both analyses, we used *F. pratensis* as outgroup to establish the ancestral state for each SNP. Some *δaδi* models were based on a non-equilibrium demography (population splits with a period of isolation before the admixture, as assumed in TREEMIX) while some were based on an equilibrium demography (fixed migration structure since divergence). We also considered the possibility of a linear population growth for some models *versus* a constant population size. In *δaδi*, we used the log L-BFGS-B optimization method to fit parameters for each model. A total of 30 independent runs were used for the optimization of each model. Each run started from a different randomly perturbed starting position and included a maximum of 20 iterations. The best diffusion fit to the observed SFS was chosen when the likelihood was the highest among the runs. Fitted models were ranked according to the Akaike information criterion (AIC) to account for the variable number of model parameters.

To set CIs for parameter estimates of the best-fit demographic models, we generated 100 datasets by non-parametric bootstrapping. Bootstrap replicates were re-optimized in *δaδi* to estimate the parameter uncertainties. CIs were calculated as *E* ± 1.96σ, where *E* is the maximum likelihood (ML) parameter estimate and σ is the standard deviation of the parameter estimate across the bootstrap replicates.

#### Origin of cultivars

We predicted the cluster membership of the 32 *L. perenne* cultivars (cultivar set) by adding them as supplementary populations in the *L. perenne* DAPC analysis of Fig. 1 (“training data”). We transformed the allele frequencies of these supplementary individuals using the centring and scaling metrics of the “training data” and determined their position onto the discriminant functions set for the “training data” (Jombart & Collins, 2015).

**Fig. 1.**
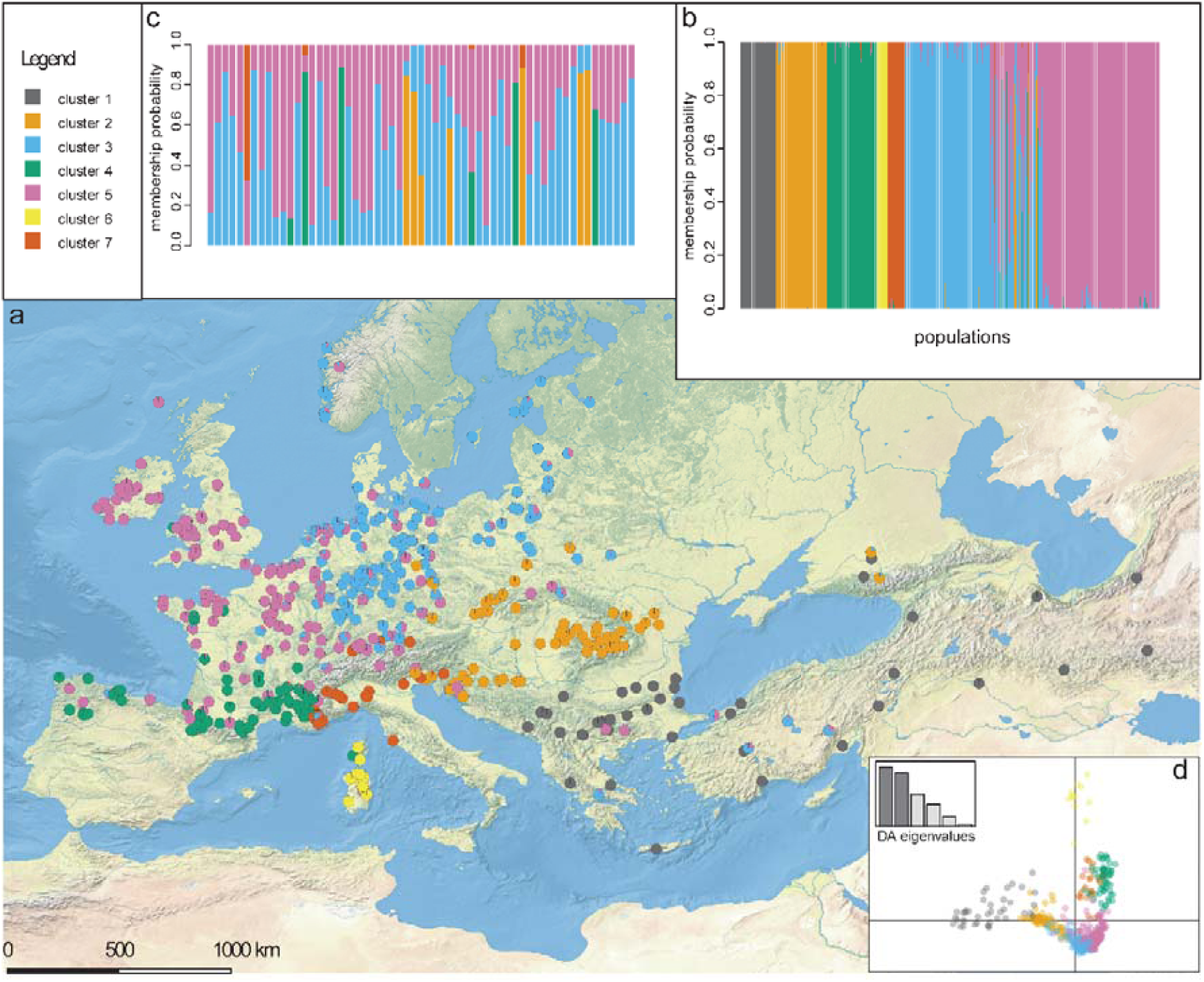
Genetic structure of 470 *L. perenne* natural populations (*L. perenne* set) sampled across Europe and the Near-East (*L. perenne set*) based on the DAPC (Discriminant Analysis of Principal Components) of 507,583 nuclear genome SNP loci. (a) Geographic distribution of genetic clusters. Pie and bar charts (b and c) indicate Populations Membership Probabilities (PMPs) to genetic clusters. (b) Representation of PMPs from all populations. (c) Zoom into PMPs of those populations having no more than 90% of membership probability in a single cluster. (d) Scatter plot of the first two discriminant functions.

## Results

### Population structure

The k-means algorithm analysis of the *L. perenne* set SNP allele frequency data found *K* = 7 as the optimal number of genetic clusters (Fig. 1). The genetic clusters exhibited a strong geographic structure and a low level of admixture (Fig. 1a). Admixture was found between clusters 3 and 5 in North-Western Europe (Fig. 1b-1c). The Discriminant Analysis of Principal Components (DAPC) axis1 was strongly correlated with longitude (r= 0.810 p-value < 0.001) and the axis2 with latitude (r= 0.662 p-value < 0.001), indicating different and independent directions of differentiation along these two geographical dimensions (Fig. S2a-S2b). Genetic differentiation among clusters was small with Fst values ranging from 0.0152 (cluster 4-cluster 5) to 0.0776 (cluster 3-cluster 6) (Supporting Information Table S1). The Mantel test indicated a significant correlation between pairwise genetic distances and geographic distances (r=0.43; p-value = 0.001) supporting the IBD hypothesis. Average expected heterozygosity (*He*) values within populations ranged from 0.54 (clusters 4) to 0.47 (cluster 6) with standard deviation (STD) within clusters ranging from 5.0E-4 (cluster 5) to 1.1E-2 (cluster 1) (Fig. 2d). Cluster 1 had a remarkably high *He* STD; this cluster indeed contains a set of populations with high heterozygosity in combination with populations with very low heterozygosity.

**Fig. 2.**
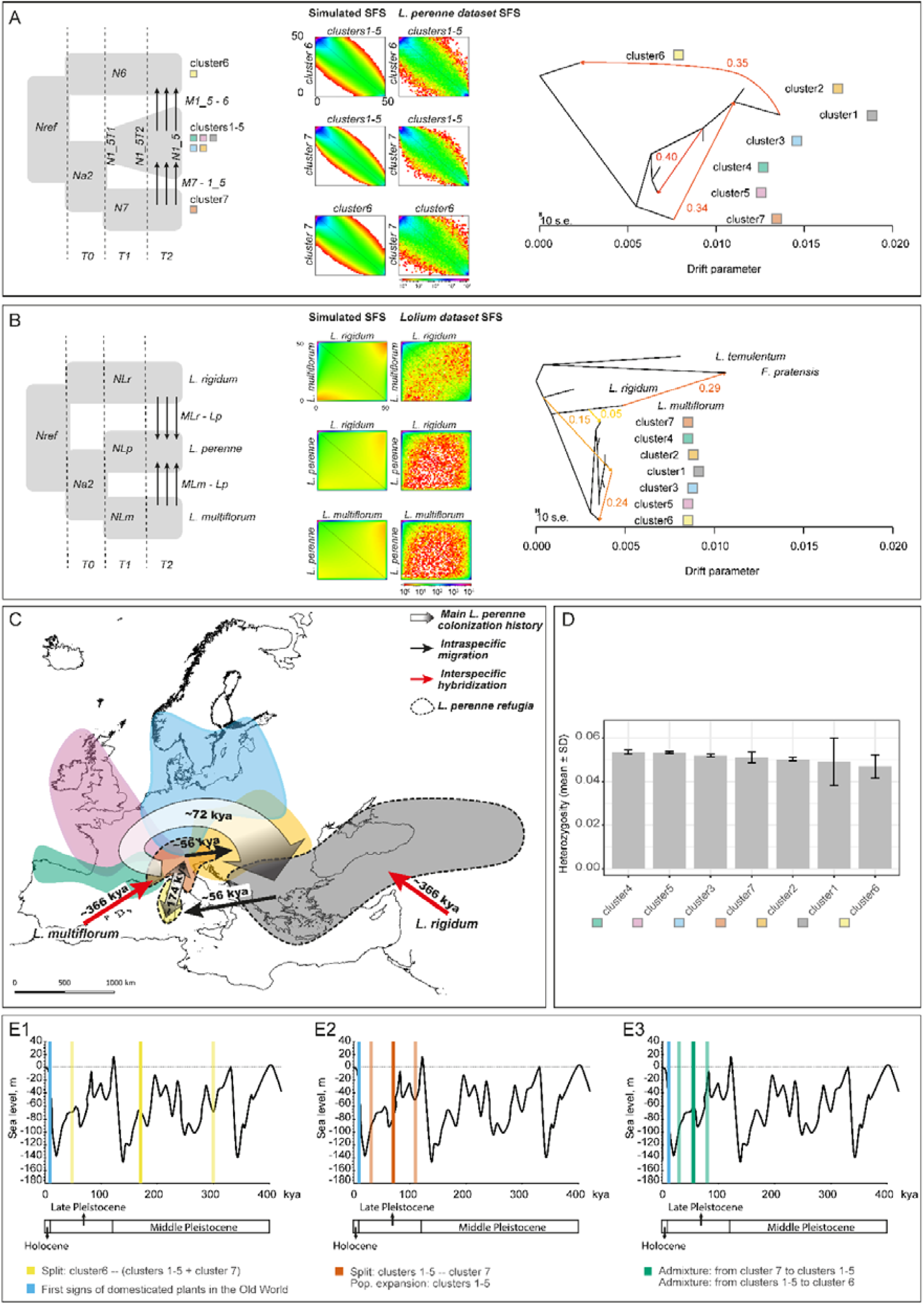
Phylogeography of *L. perenne*. (a) Analysis of *L. perenne* set only - From left to right: Schematic of the best demographic model estimated with *δaδi*, heatmap representations of the joint site frequency spectrum (SFS) expected under the best model and of the observed joint SFS of 507,583 nuclear SNPs, TREEMIX model with highest likelihood compatible with the best demographic model. (b) Analysis of *L. perenne* and related taxa populations - From left to right: Schematic of the best demographic model, heatmap representations of the joint site frequency spectrum (SFS) expected under the best model and of the observed joint SFS of 507,583 nuclear SNPs, TREEMIX model with highest likelihood compatible with the best *δaδi* model. (c) Geographic representation of the inferred demographic history of *L. perenne* in Europe. Colours represent distribution of DAPC genetic clusters as in Fig. 1, dates represent maximum likelihood values as obtained from the best *δaδi* model. (d) Mean and standard deviation of expected population heterozygosity for each *L. perenne* genetic cluster. (E-g) Simplified representation of sea level variation across the last 400 kya, from Rabineau et al. (2006) with superimposed Time parameter values: ML (opaque colours) and 95% CI (transparent colours) under the best *L. perenne δaδi* model (model F). Time estimate for the first signs of domesticated plants in the Old World *as per* (Zohary et al., 1988) is also displayed in light blue colour.

DAPC analyses carried out on 79 cpDNA loci from the *Lolium set* (*L. perenne* and other taxa) revealed neither population geographical structure in *L. perenne* nor differentiation among the different *Lolium* taxa. The K-means algorithm failed to find genetic clusters. We chose K = 4 *ad hoc* considering as optimal K the highest possible number of clusters that showed non-admixed populations. cpDNA clusters comprised members of the different *Lolium* taxa, with clusters 1 and 2 showing a higher presence in Mediterranean areas (Fig. S3).

### Splits and admixture

Setting up demographic models requires prior information about the relationships among populations. Splits and admixture analyses as implemented in TREEMIX allow to obtain the prior information necessary to set up alternative demographic models (main tree topologies and migration directions). We first analysed the *Lolium set* with TREEMIX and obtained models for increasing numbers of admixture events (M0-M6). The likelihood of the TREEMIX models (TMMs) increased almost linearly with no clear stabilization of likelihood values as the number of admixture edges increased (Fig. S4a). When adding two or more admixture edges (M2-M6) the topology of the main tree was rearranged and cluster 6 became the most basal lineage. The ln-likelihood of the *Lolium set* TMM increased by 591.37 from M0 to M1, 779.78 from M1 to M2 and 337.28 from M2 to M3. All additional edges (M4-M6 models) increased the ln-likelihood by less than 212. It is important to take into consideration that the increase of likelihood values does not mean that the model is closer to the true network. This is because the addition of admixture edges can never reduce the likelihood. Additionally, true admixture could be confounded with the effect of independently occurring mutations that are not accounted for in the TREEMIX model (Pickrell & Pritchard, 2012). All migration events between *L. multiflorum, L. rigidum* and *L. perenne* observed in alternative *Lolium set* TREEMIX models were considered for the generation of demographic models with *δaδi* (see below).

We analysed the *L. perenne set* with TREEMIX using cluster 6 as outgroup as displayed in the *Lolium set* TMMs 2-6 (Fig. S4b). With the addition of admixture edges, the likelihood of the *L. perenne* TMMs clearly increased from M0 to M2 and to a lower extent from M2 to M6 (Fig. S4b). The ln-likelihood of the *L. perenne set* TMM increased by 13,232.72 from M0 to M1, 5208.19 from M1 to M2 and 984.35 from M2 to M3. All additional edges (M4-M6 models) increased the ln-likelihood by less than 653.

### Demography

We considered all migration events observed in alternative *L. perenne* TREEMIX models for the generation of 12 *δaδi* demographic models (Fig. S5a). More specifically, we compared *δaδi* models with contrasting patterns of migration (as displayed in the alternative *L. perenne* TREEMIX models M1-M6) and population growth (linear growth vs exponential growth). Model comparison revealed that the best fit between the predicted and observed site frequency spectrum (SFS) (*i.e.* the highest maximum composite likelihood) was obtained with model F (Fig. 2a and Supporting Information Table S2). This model showed an ancestral population splitting in two lineages (cluster 6 and the ancestor of clusters 1-5 + 7), then a second split followed by a period of isolation between clusters 1-5 and cluster 7. This was followed by migration from cluster 7 to clusters 1-5 and from clusters 1-5 to cluster 6. Residuals of the model (normalized differences between model and data) followed a normal distribution with a zero mean value, indicating an appropriate fit of the model to the data. We used the best fit of the best model over 30 independent runs to get estimates of model parameters (Supporting Information Table S2). Then using a known average mutation rate of 6.03E-9 substitutions per site per generation for Poaceae (De La Torre et al., 2017), and assuming an average generation time of three years for *L. perenne* (3y/gen), we converted effective population sizes (Ni) and Time (Ti) parameters to (breeding) individuals and years (values for all models are presented in Table S3). Maximum likelihood parameter values and their 95% confidence interval (CI) obtained from non-parametric bootstrapping are shown in Supporting Information Table S4. The first ancestral population splitting into two lineages (cluster 6 and clusters 1-5 + 7) was dated 174 kya (49 – 300 kya, 95% CI). Then, a second split giving rise to the lineages containing clusters 1-5 and cluster7, followed by a linear population growth for clusters 1-5 was set at 72 kya (31 – 112 kya, 95% CI). Finally, admixture from cluster 7 to clusters 1-5 and from clusters 1-5 to cluster 6 started 56 kya (32 – 81 kya, 95% CI). Additionally, we ran an alternative to model F in *δaδi*, in which the latter two admixture events did not coincide. This new model did not fit as well as the base model F (results not shown) suggesting similar timing for the two admixture events.

Using the *Lolium set*, we compared *δaδi* models with contrasting admixture patterns between *L. rigidum, L. multiflorum* and *L. perenne* as observed with TREEMIX and also with contrasting patterns of population growth (linear growth *vs* exponential growth in *L. perenne,* Fig. S5b). Model comparison revealed that the best fit to our observed SFS was model C (Fig. 2b and Supporting Information Table S5). This model detected an ancestral population splitting into two lineages (*L. rigidum* and the ancestor of *L. perenne* and *L. multiflorum*) then a second split followed by a period of isolation between *L. perenne* and *L. multiflorum* followed by migration from *L. rigidum* to *L. perenne* and from *L. multiflorum* to *L. perenne* with constant population size in *L. perenne*. Residuals of the model showed a normal distribution with a zero mean value. To obtain parameter values, we used the best fit of model C over 30 independent runs and used the known average mutation rate of 6.03E-9 substitutions per site per generation for Poaceae (De La Torre et al., 2017). In this case, we displayed real values considering two different generation times: one year (for the annual *L. rigidum*, and the annual or short-lived perennial *L. multiflorum*; 1y/gen) and three years (for *L. perenne*; 3y/gen). Real values for all models are presented in Table S6. Maximum likelihood parameter values and their 95% CI obtained from non-parametric bootstrapping are shown in Supporting Information Table S7. The first ancestral population splitting into two lineages (*L. rigidum* and *L. perenne* + *L. multiflorum*) was dated 397 kya (238-555 kya, 1y/gen) or 1191 kya (715-167 kya, 3y/gen). A subsequent split between *L. perenne* and *L. multiflorum* occurred 380 kya (235-525 kya, 1y/gen) or 1141 kya (706-158 kya, 3y/gen). Finally admixture from *L. rigidum* to *L. perenne* and from *L. multiflorum* to *L. perenne* started 366 kya (235-497 kya, 1y/gen) or 1098 kya (704-1492 kya, 3y/gen).

### Origin of cultivars

The predicted membership of supplementary data (32 cultivars bred for forage usage during the last six decades in Europe and New Zealand) on the *L. perenne* set DAPC (Fig. 3) showed that 25 out of these 32 cultivars were assigned to cluster 5, (North-Western Europe), two to cluster 2 (Central-Eastern Europe), three to cluster 7 (Northern Italy), one to cluster 3 (North-Eastern Europe) and one to cluster 4 (South-Western Europe). Most of cultivars assigned to cluster 5 were genetically very similar to the natural populations of this cluster. Cultivars assigned to the other clusters were highly admixed, with high membership probabilities with cluster 5 except the cultivar *Medea*. A DAPC analysis with all populations did not separate cultivars from natural populations (not shown). This suggests that the genetic origin of cultivars is not restricted to a single cluster, despite the major contribution of cluster 5.

**Fig. 3.**
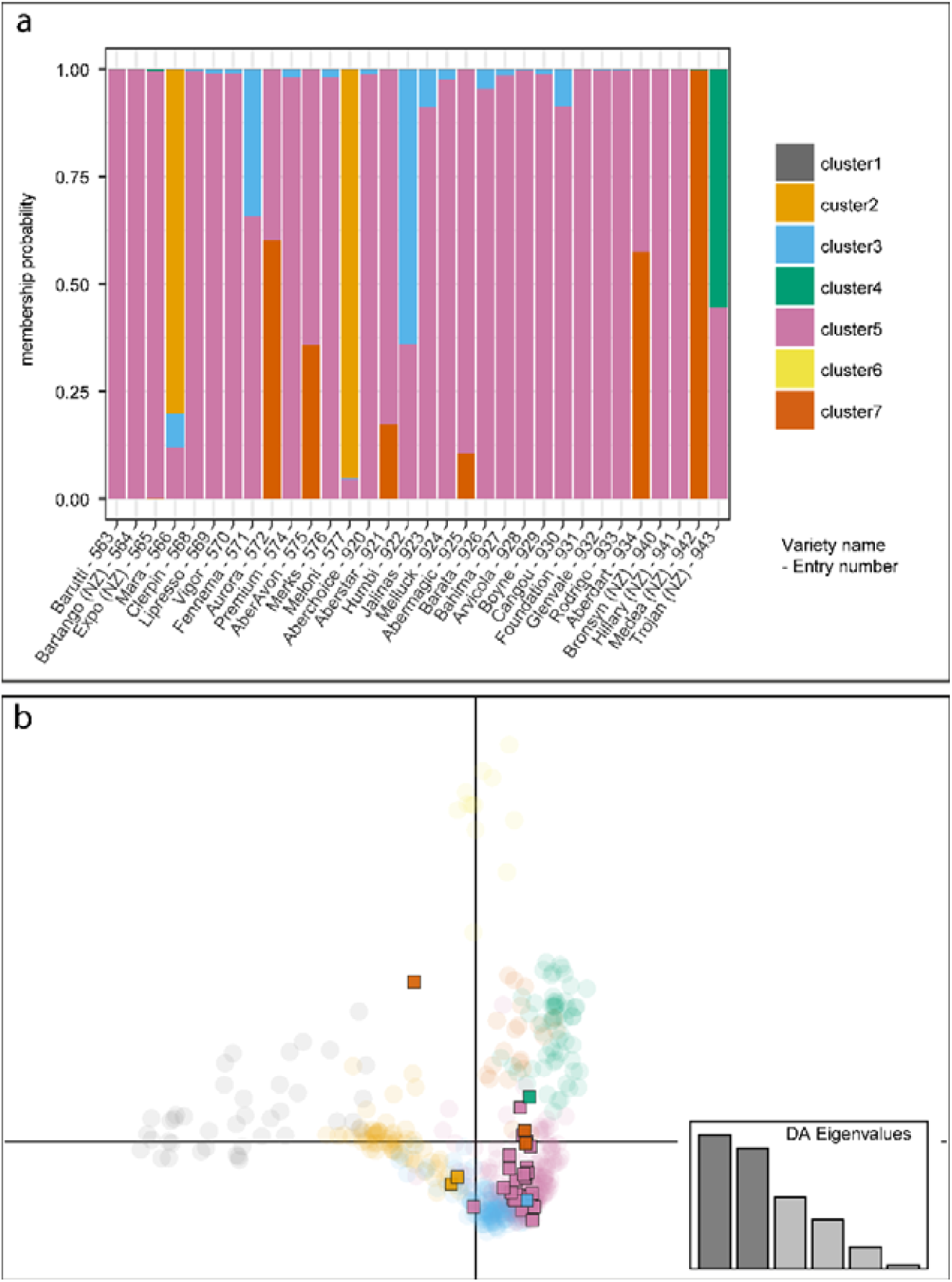
Prediction of DAPC (Discriminant Analysis of Principal Components) cluster membership for a set of 32 *L. perenne* cultivars bred for forage usage and released within the six last decades in Europe and New Zealand. (a) Membership probability with respect to a DAPC constructed with 470 *L. perenne* natural populations. (b) Scatter plot of the first two discriminant functions of the DAPC with *L. perenne* natural populations displayed as transparent filled circles (same positions as on Fig. 1) and cultivars superimposed as opaque filled squares.

## Discussion

According to the fossil record, the Late Glacial (ca. 13-10 kya) vegetation in Europe was dominated by herbaceous communities including a large proportion of grasses (Giesecke et al., 2017). Later on, during the Holocene (from ca. 10 kya onwards), and particularly during the Holocene Climatic Optimum, Europe became dominated by dense forests of temperate deciduous trees and conifers (Feurdean et al., 2015; Giesecke et al., 2017). But forests were never fully closed, enabling the persistence of grasslands throughout the Holocene (Hejcman et al., 2013). For example, small-scale steppe grasslands of natural origin were present in forest-rich areas of Central Europe before the onset of agriculture in the early Neolithic (ca. 5.5 kya) (Hejcman et al., 2013). In the late Holocene (last 3.5 kya), an increase in the abundance of grasses, accompanied by a reduction of forested areas is captured in the fossil record, reflecting the development and expansion of agriculture in this area of Europe (Hejcman et al., 2013; Giesecke et al., 2017). Up to date, the impact of this anthropogenic transformation of the European landscapes on the genetic diversity of key grassland species has been scarcely documented.

The processes involved in the genesis of grasslands in agricultural landscapes are not precisely known. The unconscious selection of grazing-tolerant species has certainly played a major role and has been pivotal in the domestication of large herbivores. However, the relevance of conscious human actions (such as voluntary seed harvesting and sowing of grasslands) remains largely unknown, and no fossil record specific for *L. perenne* or other major grass species has been reported at documented archaeological sites. Nonetheless, we can ascertain that intensive grassland cultivation may not have taken place before the appearance of the first scythes during the 7th– 6th century BC (Hejcman et al., 2013) and that the development of specific practices for the management and production of grasslands occurred during The Roman Empire (Hooper & Ash, 1935; Ash et al., 1941). Our analyses of the genomic diversity, structure, and past demography of *L. perenne* reveal that more than two and a half millennia of extensive use did not have a notable impact on the diversity of extant wild populations of this dominant grass species. This is supported by the clear geographical structure found in this species (Fig. 1) and by the estimated time when main demographic events took place (Fig. 2c and Supporting Information Table S4).

The origin of *L. perenne* and its close relatives (*L. rigidum* and *L. multiflorum*) inferred by our analyses is congruent with previous studies (Balfourier et al., 1998; Catalán et al., 2004). This includes the early divergence of *L. rigidum* and the sister relationship between *L. multiflorum* and *L. perenne*. However, we discovered that relationships between these three taxa are more complex than previously proposed with *L. perenne* receiving genes from *L. rigidum* and *L. multiflorum* after divergence (Fig. 2b-2c). According to our demographic reconstructions, main events took place during the Pleistocene glaciations, long before the onset of agriculture and main migrations of agricultural communities across Europe. This is in agreement with previous inferences from molecular dating of the grass genera *Festuca* and *Lolium* (Inda et al., 2014). Gene flow from *L. rigidum* and *L. multiflorum* to *L. perenne* (Fig. 2b-2c) started 366 kya (1y/gen) or 1190 kya (3y/gen) (ML values, see Supporting Information Table S7). A close relationship between Middle-Eastern ancestral *L. perenne* populations and *L. rigidum* was previously inferred (Balfourier et al., 1998). Also, *L. multiflorum* was acknowledged as being present in the Italian plains since the High Middle Ages, but it was wrongly assumed that it originated from human-mediated selection of *L. perenne* genotypes from this region (Casler, 2006). It is thus not surprising to find genetic material from *L. rigidum* and *L. multiflorum* into *L. perenne* populations from Middle-East (cluster 1) and Italy (cluster 7), respectively (see Fig. S4a). The next event reconstructed from the best *L. perenne* model (Fig. 2a) is a split between cluster 6 and the ancestor of remaining clusters occurring 174 kya (Fig. 2c and Supporting Information Table S4). Despite the broad CI (Fig. 2e and Supporting Information Table S4), this split overlaps with a period of time with important sea level variation in the Mediterranean basin (Rabineau et al., 2006) (Fig. 2e). A sea level rise might have caused the isolation between cluster 6 in Corsica and Sardinia and Italian refuge lineages already present there (as shown by the ancestral admixture signal of *L. multiflorum*, Fig. 2b, Fig. 2c and Supporting Information Table S7). The Corsica channel might have acted as a barrier for gene flow in *L. perenne*, as previously reported for other Mediterranean species such as *Cistus creticus* (Falchi et al., 2009). The next event is the split between cluster 7 and clusters 1-5, followed by expansion of clusters 1-5 towards South-Western and North-Western Europe and next towards Eastern Europe (clockwise movement around the Alps) which is dated 72 kya (Fig. 2a, Fig. 2c and Supporting Information Table S4). These events overlap with a glacial period (Würm glaciation 12-110 kya) (Fig. 2f). During the Würm glaciation, the Alps might have acted as a barrier to gene flow in *L. perenne* as previously suggested by Balfourier et *al.* (Balfourier et al., 1998, 2000) and Blackmore et *al.* (Blackmore et al., 2015). In continental Europe, the continuous cooling during that period seems to have affected the dominance of tree species in favour of herbs, including grass species such as *L. perenne*. This is in agreement with the abundance estimates traced in the pollen record for different vegetation types during the Late Glacial (Giesecke et al., 2017). During the eastward expansion of *L. perenne*, the leading edge of clusters 1-5 may have contacted ancient *L. perenne* genotypes already located in Eastern Mediterranean areas which had previously admixed with *L. rigidum* (cluster 1, see Fig. 2c). This is further supported by the analysis of genetic diversity (*He*) of *L. perenne* clusters (Fig. 2d) which showed that cluster 1 exhibits the most variable within-population genetic diversity. Cluster 1 may have been formed by the mixture between ancient highly diverse populations located in South-Eastern refugia and more recent immigrant populations from cluster 2, the latter including low genetic diversity due to allele surfing in expanding populations (Edmonds et al., 2004; Excoffier & Ray, 2008). In later stages of the Würm glaciation from 56 kya until current time, the best demographic model suggest migration from cluster 7 to clusters 1-2 and from cluster 1 to cluster 6 (Fig. 2a, Fig. 2c and Supporting Information Table S4). During this period, and particularly 20 kya, the Adriatic sea level reached an estimated maximum drop of 149 meters with respect to current level (Lambeck, 2001), which may have facilitated dispersal. Variation in the Mediterranean sea level was previously invoked to explain plant species colonization routes inferred from genetic data (Petit et al., 2002; Fernández-Mazuecos & Vargas, 2011; Lo Presti & Oberprieler, 2011; Garnatje et al., 2013). In this case the Adriatic Sea level drop may have narrowed straits, thus facilitating dispersal events that led to colonization and admixture among *L. perenne* lineages.

Since the origin and expansion of domesticated plants in the Old World, trade, wars and nomadism caused extensive movements of people and livestock across Europe, possibly favouring transport of diaspores of grassland species among European regions. Nevertheless, all expansion and admixture events in *L. perenne* recovered by our *δaδi* analyses predate the origin of agriculture in Europe, even when taking into account the CI of our time estimates (see Fig. 2e-2g and Supporting Information Table S4 and Table S7). The results we report reveal that main genetic signals of colonization and admixture observed in *L. perenne*, a dominant grass species in European grasslands, can be explained by climatic events predating the transformation of landscapes by human activities in Europe. This study provides the clearest evidence of a limited effect of the agricultural expansion in the genetic diversity of European grasslands. A much more limited dataset previously reported similar conclusions for *F. pratensis* (Fjellheim et al., 2006). These two major grass species of European grasslands (*L. perenne* and *F. pratensis*) may have thus experienced similar evolutionary histories. They may therefore exemplify an evolutionary trend shared by most European grasslands species, in which the current genetic diversity patterns were shaped during range expansions of the last glacial period.

The first detailed written records on intensification of grassland management date from The Roman Empire (Hooper & Ash, 1935; Ash et al., 1941). However, the real decline of wild grasslands and the large-scale enlargement of hay meadows started in many European regions during the 18th century (Semelová et al., 2008). This process may have involved increased use of grass seeds harvested in local natural strains to re-sow pastures, but likely without direct selection on phenotypic traits except sufficient seed production. Modern selection of *L. perenne* for forage production based on the agronomic testing of progeny performance began in 1919 in the United Kingdom and after World War II in other European countries, Northern America and New Zealand (Humphreys et al., 2010; Sampoux et al., 2011). Recurrent selection in plant breeding germplasm has so far resulted in the release of more than 1000 *L. perenne* cultivars improved for forage production (European Commission, 2017). More recently, since years 1960, similar selection methods have also been implemented to release *L. perenne* cultivars improved for turf usage (Sampoux et al., 2013). We genotyped 32 cultivars representing a large diversity of the *L. perenne* cultivars bred for forage usage in various countries of Europe and New Zealand. 25 out of the 32 genotyped cultivars were assigned to the cluster of North-Western Europe populations and were thus likely bred from germplasm of this area. Cultivars assigned to other clusters, except Medea, also contained a significant part of diversity from the North-Western Europe cluster 5. An interesting pattern is observed in a set of cultivars developed by IBERS (Aberystwyth, UK): Aurora, a cultivar with high water soluble carbohydrate (WSC) content developed from a Swiss ecotype (Faville et al., 2004) showed 60 % membership of cluster 7. Conversely, other more recent IBERS cultivars developed from Aurora (Aberdart, Aberavon, Aberstar and Abermagic) showed a decreasing content of genetic material from cluster 7 in favour of cluster 5. This case exemplifies the strong bias towards North-Western Europe cluster 5 in the generation of modern cultivars. Most probably, this trend is predominantly due to the fact that modern breeding of *L. perenne* started in this part of the world using local diversity.

We show that *L. perenne* cultivars likely use only a small proportion of the natural genetic diversity existing across its natural distribution range. This natural diversity thus represents a much valuable genetic resource that should be safeguarded in genebanks, but also much more efficiently in the diverse natural grasslands from which it originates. However, since the mid-20^th^ century, there has been a continuous trend of reduction in the acreage of natural grasslands in Europe. Especially in intensive agricultural landscapes, natural grasslands have tended to be ploughed and replaced by rotations of annual crops. Rotations may indeed include temporary meadows sown with cultivars from modern breeding in regions where agriculture combines cash crops and cattle breeding. This practice may not only reduce natural diversity of perennial ryegrass by extinction of natural populations but also by the expansion of North-Western European genotypes (typically found in cultivars) into other European regions through gene flow from cultivars to natural populations.

Demographic reconstructions assessing the SFS from Pool-seq GBS performed on a high number of populations allowed to trace back the origins of the genomic diversity of *L. perenne* at an unprecedented level of detail. The current extent of grasslands across Europe has been mainly determined by human activities. However, our results indicate that the spatial distribution of the natural genome-wide diversity of *L. perenne* has not been significantly disturbed after more than two millennia of agricultural expansion. The current *L. perenne* natural populations still maintain the genomic diversity that has allowed the species to persist during the Quaternary climatic fluctuations. Modern plant breeding has likely used only a small part of the genomic diversity of *L. perenne* naturally distributed across and around Europe, and thus has likely taken limited advantage of the adaptive diversity and phenotypic variability of the species. To date, this wide natural diversity remains available but it is endangered and should be preserved. Indeed it may constitute a much valuable genetic resource for plant breeding to meet emerging agricultural challenges and more specifically to adapt *L. perenne* to the anthropogenic climate change foreseen to occur in the next decades.

## Biosketch

José Luis Blanco Pastor’ s main scientific goal is to study *the evolution and diversity of species, populations and genes*. Specifically his research is focussed on the study of the effect of past climate changes on plant evolution, gene flow and adaptation to environmental factors. He is particularly interested in the use of this information to help plant species to adapt to future climatic conditions. JLBP, SM, PB, EW, KJD, MH, TR, IRR and JPS designed research; JLBP analysed data; JPS, AMR, EW, KJD, MH, HM, TR and TL collected data; JLBP, SM, PB, AEG and JPS interpreted results; JLBP and JPS wrote the manuscript with feedback from SM, TR and IRR. All authors participated in the edition of the manuscript.

## Data accessibility Statement

The genetic data reported in this study are available in the NCBI Short Read Archive (SRA) database through accession SRP136600.

## Supporting Information Legends

Fig. S1. Value of BIC for each number K of clusters as obtained with the K-means algorithm implemented in adegenet Jombart, et al. (2010) applied to the Lolium perenne set.

Fig. S2. Geographical pattern of genetic differentiation in *L. perenne*. Scatterplot of longitude vs principal component 1 and scatterplot of latitude vs principal component 2.

Fig. S3. Genetic structure of the *Lolium set* based on the DAPC analysis of 79 cpDNA SNPs. Fig. S4. *Lolium set* and *L. perenne set* TREEMIX models.

Fig. S5. Schematic representation of *L. perenne* set and *Lolium set* δaδi models.

Fig. S6. 50% Majority-rule consensus tree of 552 accessions of Lolium perenne and related taxa.

Fig. S7. Scatterplot of the first two principal axes from a PCA of 552 accessions of *Lolium perenne* and related taxa computed using allele frequencies of 507,583 nuclear SNPs.

Table S1. Fst statistics between seven clusters discriminated by a DAPC (Discriminant Analysis of Principal Components) of the *L. perenne set.*

Table S2. Parameter values from the fitting of 12 alternative *∂*a*∂*i demographic models using the *L. perenne set.*

Table S3. Parameter values from the fitting of 12 alternative *∂*a*∂*i demographic models using the *L. perenne se*t with *Ni* and *Ti* values converted to numbers of individuals and years, respectively.

Table S4. Maximum Likelihood parameter values and non-parametric bootstrap 95% confidence interval of the best *L. perenne set* demographic model.

Table S5. Parameter values from the fitting of 12 alternative *∂*a*∂*i interspecific models of gene flow using the *Lolium set.*

Table S6. Parameter values from the fitting of 12 alternative *∂*a*∂*i interspecific models of gene flow using the *Lolium se*t with *Ni* and *Ti* values converted to numbers of individuals and years, respectively.

Table S7. Maximum Likelihood parameter values and non-parametric bootstrap 95% confidence interval of the best *Lolium* set gene flow model.

Table S8. Accessions from *Lolium perenne* and related taxa used in the study.

Table S9. cpDNA HiPlex Primer sequences.

Methods S1. Plant Material and Genotyping.

## Acknowledgements

J.L. Blanco-Pastor has received the support of the EU in the framework of the Marie-Curie FP7 COFUND People Program, through the award of an AgreenSkills+ fellowship (under grant agreement n° 609398). This work was funded in the frame of the project *GrassLandscape* awarded by the 2014 FACCE-JPI ERA-NET+ call *Climate Smart Agriculture*. Funding was granted by the European Commission (EC grant agreement n° 618105), by the Agence Nationale de la Recherche (ANR) and the Institut National de la Recherche Agronomique (INRA – métaprogramme ACCAF) in France, the Biotechnology and Biological Sciences Research Council (BBSRC) in the United-Kingdom, the Bundesantalt für Landwirtschaft und Ernährung (BLE) in Germany.

The authors thank the curators from the genebanks which contributed perennial ryegrass seed samples for the needs of the project. Perennial ryegrass is one of the plant species covered under the Multilateral System of the International Treaty on Plant Genetic Resources for Food and Agriculture. All genetic materials used in this study were made available to the authors after signature of a Standard Material Transfer Agreement (SMTA) by the provider and the recipient. Implementation and signature of a SMTA provides compliance with the provisions of the Nagoya Protocol for parties wishing to provide and receive genetic material under the Multilateral System. The authors declare no conflict of interest.

